# MiRNA let-7a-5p Ameliorates Pulmonary Fibrosis by Suppressing TGFBR1-Mediated Endothelial-to-Mesenchymal Transition

**DOI:** 10.64898/2026.08.18.745407

**Authors:** Juan Pang, Jiarui Shen, Wei Yang, Zhihui Wu, Xing Gu, Yuhang Xia, Ruixuan Wang, Longzhi Wang, Yujie Cao, Jianying Li, Hui Shen, Fenqing Shang

## Abstract

**Background:** Idiopathic Pulmonary Fibrosis (IPF) is a fatal chronic lung disease with limited therapeutic options. While alveolar epithelial injury and fibroblast activation are well-studied, endothelial-mesenchymal transition (EndoMT) is emerging as a critical pathogenic mechanism. The regulatory role of exosomal miRNAs in pulmonary fibrosis remains unclear. This study investigates serum exosomal miRNAs, particularly let-7a-5p, in modulating EndoMT during the onset of pulmonary fibrosis.

**Methods:** Clinical cohorts of IPF patients and healthy controls were enrolled. Serum exosomal miRNAs were profiled, followed by differential expression and functional enrichment analyses. In vitro experiments involved human pulmonary artery endothelial cells (HPAECs) transfected with let-7a-5p mimic or inhibitor. Dual-luciferase reporter assays confirmed the binding between let-7a-5p and TGFBR1. HPAECs were co-cultured with lung epithelial cells to examine paracrine signaling. In vivo studies used a bleomycin-induced mouse model with let-7a-5p agomir administration. Assessments included histopathological staining, hydroxyproline content, Western blot, qPCR, micro-CT, and pulmonary function tests.

**Results:** Let-7a-5p was significantly downregulated in serum exosomes from IPF patients, correlating with clinical indicators. Mechanistically, let-7a-5p directly bound the TGFBR1 3′UTR to inhibit its expression. Inhibition of let-7a-5p upregulated α-SMA, FN1, smad2/3 phosphorylation, and collagen I, while downregulating CD31 and VE-cadherin. Therapeutically, let-7a-5p mimic reversed bleomycin-induced EndoMT and suppressed epithelial-mesenchymal transition (EMT) via paracrine signaling. Mice administered agomir showed reduced fibrosis, improved lung function, and suppressed TGF-β/Smad signaling.

**Conclusion:** Serum exosomal let-7a-5p suppresses pulmonary fibrosis by targeting TGFBR1 to inhibit EndoMT. Its downregulation in IPF patients correlates with disease progression, highlighting its biomarker potential.

## INTRODUCTION

Idiopathic pulmonary fibrosis is a severe chronic lung disease characterized by progressive scarring of lung tissue^[1, 2]^. As such, the normal structure of the lung interstitium is destroyed, leading to impaired effective expansion and contraction of the lungs, which results in dyspnea in patients^[3]^. Currently, clinical treatment options, such as pirfenidone and nintedanib, are due to their anti-fibrotic anti-oxidative, and anti-inflammatory effects^[4, 5]^. However, these agents can only slow the progression of the disease but cannot reverse fibrosis^[6]^. Therefore, there is unmet need to explore new pathogenic mechanisms and to seek effective therapeutic targets.

Current investigations on IPF predominantly focus on the injury of alveolar epithelial cells and activation of fibroblasts, with epithelial-mesenchymal transition (EMT) being the principal mechanism^[7–9]^. Nevertheless, the role of vascular endothelial cells (ECs) in the pathogenesis and advancement of IPF has gained attention^[10, 11]^. A study involving lineage tracing indicated that approximately 16% of pulmonary fibroblasts in bleomycin-treated mice originated from ECs^[12]^. Endothelial-mesenchymal transition (EndoMT) is evident in bleomycin and silica-induced pulmonary fibrosis in mice^[13, 14]^. With pulmonary fibrosis presented in 50-80% of patients with systemic sclerosis, cell undergoing EndoMT have been detected in the cutaneous vasculature of these patients^[15, 16]^. The derived myofibroblasts secrete ECM, thus aggravating the fibrotic lesions. Moreover, the EndoMT process may affect ECM degradation and remodeling, thereby modulating the progression of pulmonary fibrosis^[11, 17]^. EndoMT involves multiple pathways, including TGF-β/Smad, BMP, and Notch signaling^[18–21]^.

Exosomes are extracellular vesicles with diameters ranging from 30 to 150 nm, which exert endocrine and paracrine effects in health and disease, and thus providing new opportunity for drug delivery and targeting^[22–24]^. With respect to pulmonary fibrosis, it remains ambiguous how exosomal mediates small RNAs to selectively sorted and secreted into the circulation, and how they regulate EndoMT. The let-7 miRNA family is widely involved in physiology and various diseases. It has been suggested that let-7 acts as a central hub linking impaired plasticity in type II alveolar cells with fibrogenesis^[25]^. Let-7a-5p exhibits anti-inflammatory and antiviral properties, playing a key role in acute lung injury^[26, 27]^. Diazepam delays pulmonary fibrosis progression by suppressing the let-7a-5p/MYD88-mediated pyroptosis and inflammation^[28]^. Previous studies on the let-7 family in relation to pulmonary fibrosis have mostly focused on EMT, with relatively few investigations into its association with EndoMT, and almost no studies on let-7a-5p specifically in the context of EndoMT. Given the pathogenic role of EndoMT in pulmonary fibrosis, we report herein that in human IPF and mouse models of pulmonary fibrosis, let-7a-5p can specifically target TGFBR1 mRNA, thereby suppressing EndoMT in lung endothelial cells and also influencing EMT through paracrine signaling. This study provides novel evidence for the anti-fibrotic mechanism of the let-7 family, specifically demonstrating its role in endothelial-mesenchymal transition (EndoMT).

## MATERIALS AND METHODS

### Serum and Clinical Data in Pulmonary Fibrosis

This retrospective study utilized archived serum samples and clinical data from patients diagnosed with pulmonary fibrosis at Xi’an Chest Hospital between September 2023 and November 2025. All patients were diagnosed based on clinical and imaging examinations according to the 2016 Chinese Expert Consensus on the Diagnosis and Treatment of Idiopathic Pulmonary Fibrosis. Serum samples from matched healthy volunteers were collected as controls. Clinical parameters were systematically collected from electronic medical records. The data were accessed for research purposes between December 2025 and January 2026, during which all analyses were completed. All patient records and serum sample data were de-identified and anonymized prior to the authors’ access; therefore, the authors did not have access to information that could identify individual participants during or after data collection. The study was approved by the Ethics Committee of Xi’an Chest Hospital (Approval No. R2025-012-01), and the requirement for informed consent was waived due to the retrospective nature of the study and the use of anonymized data. A radiologist scored CT images using the Camiciottoli visual scoring method, which evaluates lesion type and lesion extent. Lesion type scores: ground-glass opacity (1 point), irregular pleural border (2 points), irregular interlobular septal thickening (3 points), honeycombing (4 points), and subpleural bullae (5 points). Lesion extent scores: involvement of 1–3 lung segments (1 point), 4–9 lung segments (2 points), and >9 lung segments (3 points). The total score (range 0–30) was the sum of lesion type and extent scores (a score of 0 was assigned if none of the five lesion types were present).

### Serum KL-6 Detection

Serum KL-6 levels were measured using an ELISA kit (Shanghai Enzyme-linked Biotechnology Co., Ltd.; ml057455A), involving incubation of diluted samples and standards, sequential addition of a biotinylated anti-KL-6 antibody and SA-HRP conjugate, follow by chromogen development, and absorbance reading at 450 nm, with concentrations determined via standard curve interpolation.

### MiRNA Sequencing and Data Analysis

Total RNA was isolated from whole blood using the PAXgene® Blood RNA Kit (Qiagen, 762174); RNA purity was assessed using a NanoDrop 2000 spectrophotometer (Thermo Scientific) and integrity confirmed via Agilent 2100 Bioanalyzer (Agilent Technologies). High-quality RNA was used to construct small RNA sequencing libraries following the protocol of the VAHTS Small RNA Library Prep Kit for Illumina V2 (Vazyme, NR811), including 3’/5’ adapter ligation, reverse transcription, and PCR enrichment. Libraries were sequenced as single-end 50-bp reads on an Illumina HiSeq 2000. Raw reads were trimmed and filtered with Trimmomatic v0.30, removing adapters, low-quality bases (Q < 20), “N”-containing ends, sequences with average Q < 20 in a 4-bp sliding window, and reads shorter than 18 bp post-trimming. Clean reads were aligned and annotated against the human genome using miRDeep2, retaining only miRNAs with a mean read count > 2 across the eight samples. Differential expression analysis was performed with DESeq2 v1.46.0, identifying significant miRNAs at P < 0.05 and |log₂ (fold change) | > 1. Target prediction employed TargetScanHuman 8.0, regulatory networks were visualized in Cytoscape v3.10, and functional enrichment (GO/KEGG) was conducted using clusterProfiler v4.16.4, with statistical significance determined by hypergeometric testing.

### Cell Culture

Human pulmonary artery endothelial cells (HPAECs, Shanghai Jinyuan Biotechnology) were cultured in complete medium (Jinyuan, JY-Y652) at 37°C, 5% CO₂. Cells at ∼80% confluence in 6-well plates (1×10⁶ cells/well) were transfected using lipo2000 (ThermoFisher) with let-7a-5p mimic/inhibitor or controls (Shanghai GenePharma), followed by 25 μg/mL bleomycin treatment at 6h post-transfection. For immunofluorescence, cells fixed in 4% paraformaldehyde were permeabilized (except for CD31), blocked with goat serum, incubated overnight with primary antibody anti-α-SMA (CST, 48938) and fluorescent secondary antibody, then DAPI-stained and imaged using ImageJ. For dual-luciferase assays, 293T cells were co-transfected with let-7a-5p mimic/inhibitor (30 pmol) and wild-type or mutant TGFBR1 plasmids into GP-miRGLO vector (Shanghai GenePharma) using lipo2000. Luciferase activity was measured 24h post-transfection using the Dual-Luciferase Reporter Assay Kit (Shanghai GenePharma), with firefly luciferase quantified first followed by Renilla luciferase to assess let-7a-5p-TGFBR1 binding activity. The sequences of let-7a-5p mimic, inhibitor, and their respective negative controls are shown in Supplementary Table S1. The sequences of TGFBR1 wild-type and mutant plasmids are shown in Supplementary Table S2

### Western Blot and Polymerase Chain Reaction (PCR)

Cells or tissues were subjected to lysis, and the supernatants were harvested after centrifugation. The protein concentration was ascertained using a BCA assay. Equivalent quantities of protein were resolved by SDS-PAGE using protein markers (Thermo Fisher Scientific, 26619; UElandy, P8028M; GenScript, M00521), transferred onto membranes, and incubated with primary and secondary antibodies. Signals were detected using enhanced chemiluminescence substrate (Servicebio, G2014 or Vazyme, E423-01) and exposed to X-ray film (FUJIFILM) followed by development with developer (Solarbio, YA0370) and fixer (Solarbio, YA0380), or visualized using a chemiluminescence imaging system. The images were analyzed with ImageJ software. Antibodies employed were as follows: CD31 (ABclonal, Cat: A19014), VE-cadherin (ABclonal, A12416), α-SMA (CST, 19245), Collagen I (CST, 72026), FN1 (Abcam, ab2413), TGFBR1 (CST, 49728S), smad2/3 (CST, 3102S), p-smad2/3 (CST, 8828).

RNA was isolated from cells or tissues via the Trizol method, followed by purification with chloroform, isopropanol, and 75% ethanol. For mRNA analysis, an mRNA reverse transcription kit (Vazyme, R222-01) and an amplification kit (GenScript, A308-05) were employed. For miRNA analysis, a miRNA reverse transcription kit (Thermo, 4366596) and an amplification kit (Promega, A6001) were utilized. Exosomal miRNA were isolated from serum via the exoEasy Maxi Kit (QIAGEN, 77144). The primer sequences for RNA are provided in Table S3 of the supplementary materials.

### Animal Treatment

To establish pulmonary fibrosis models, male C57BL/6J mice (6-8 weeks, 20-22 g; Beijing Vital River Laboratory Animal Technology Co., Ltd.) received a single intratracheal instillation of bleomycin (5 mg/kg; Shanghai Macklin Biochemical Co., Ltd.) on day 0, with tissue collection on day 21. Anesthesia was induced with 3–5% isoflurane and maintained with 1.5–2% isoflurane in 100% oxygen; adequate anesthesia was confirmed by absence of pedal withdrawal reflex. Mice were housed in groups of five per cage under standard conditions: temperature 22 ± 2°C, humidity 50 ± 10%, and a 12-hour light/dark cycle, with ad libitum access to food and water. Sample size was based on previous experience and field conventions. For in vivo transfection, thirty mice were divided into three groups (n=10 each): (1) control (saline intratracheally on day 0 + agomir NC twice-weekly via tail vein from day 1); (2) agomir NC + BLM (bleomycin on day 0 + agomir NC twice-weekly from day 1); (3) agomir + BLM (bleomycin on day 0 + let-7a-5p agomir (15 mg/kg; Shanghai GenePharma) twice-weekly from day 1). Body weight was monitored biweekly. All mice were euthanized on day 21 by isoflurane overdose (5% isoflurane maintained for >5 minutes after cessation of spontaneous breathing) followed by cervical dislocation. All animal procedures complied with the AVMA Guidelines for the Euthanasia of Animals.

### Histopathological Analysis

Histopathological analysis involved Hematoxylin-eosin (HE) staining, immunohistochemistry (IHC), Masson staining, and pathological scoring for alveolitis and pulmonary fibrosis. Lung tissues were fixed in 4% paraformaldehyde, paraffin-embedded, and sectioned. HE staining followed standard protocols including hematoxylin and eosin steps. Alveolitis severity was graded (0-3) using the Szapiel scoring system based on mononuclear cell infiltration and affected lung area. Pulmonary fibrosis was assessed (grades 0-8) via the Ashcroft scale, evaluating fibrotic changes and structural distortion. IHC required antigen retrieval, blocking, and incubation with primary and secondary antibodies before DAB staining and hematoxylin counterstaining. Masson staining (Beyotime, C0189S) visualized collagen with hematoxylin, Ponceau-acid fuchsin, and light green. All slides were digitally scanned (ZEISS ZEN 3.9), and quantitative analysis of IHC-positive staining and Masson collagen deposition area was performed using ImageJ.

### Measurement of Hydroxyproline Content

The content of hydroxyproline was determined using a hydroxyproline assay kit (Solarbio, QS1907). Lung tissues were hydrolyzed according to the manufacturer’s instructions, and the absorbance at 550 nm was measured to compute the hydroxyproline concentration.

### Pulmonary Function

Pulmonary function was evaluated using a BUXCO non-invasive pulmonary function system (Data Sciences International, USA). Mice were acclimatized in a dark chamber for 3 min until their breathing reached a stable state. Respiratory parameters were recorded over a period of 5 min. The following parameters were obtained: tidal volume (TVb), minute ventilation (MVb), peak expiratory flow (PEF), mid - expiratory flow (EF50), peak expiratory time ratio (Rpef), and relaxation time (Tr). The mean value of each parameter was utilized for analysis.

### Micro-Computed Tomography (Micro-CT)

Mice were anesthetized with 1.5% isoflurane and subjected to scanning using a Scanco VivaCT80 system (Scanco Medical AG, Switzerland). The scanning parameters were as follows: resolution = 30 μm, exposure time = 300 ms, voltage = 70 kV, current = 114 μA, and scan time = 8 min. Images from the same anatomical level were compared among groups. High-density white patches were indicative of fibrotic regions.

### Statistical Analysis

Statistical analyses were performed using GraphPad Prism 9.0.0 (GraphPad Software, San Diego, CA, USA). Data are presented as mean ± SEM. The Shapiro-Wilk test was used to assess normality of data distribution. For comparisons between two groups, the unpaired t-test (for normally distributed data) or the Mann-Whitney U test (for non-normally distributed data) was used. For multi-group comparisons, one-way ANOVA followed by Tukey’s post hoc test (for normally distributed data) or the Kruskal-Wallis test followed by Dunn’s post hoc test (for non-normally distributed data) was used. Body weight changes over time were analyzed using two-way repeated measures ANOVA with Bonferroni’s post hoc test. Correlations between variables were assessed using Pearson’s (parametric) or Spearman’s rank (non-parametric) correlation coefficient, depending on data distribution. Statistical significance was set at p < 0.05.

## RESULTS

### Let-7a-5p is decreased in IPF patients

Sera were procured from patients diagnosed with IPF and healthy control subjects. Subsequently, the serum exosomal miRNAs were isolated. Among these miRNAs, the expression of let-7a-5p was notably downregulated (Figure 1A-1B). Gene function enrichment analysis of the miRNA-targeted genes unveiled associations with mesenchymal transformation and the TGF-β pathway (Figure 1C). The intersection between putative miRNAs that upregulated genes in the lung tissues of IPF patients retrieved from the public database GSE53845 and the sequencing results of the present study led to 37 overlapping miRNAs (Figure 1D-1E). A total of 10 miRNAs with relatively higher abundance and a significant p-value of 0.01 were chosen for validation. For this, the sample size was expanded to 10 cases per group for PCR validation. The results showed that the level of 10 miRNAs, including let-7a-5p, let-7c-5p, and let-7e-5p, was reduced, consistent with the sequencing results (Figure 1F).Among the serum exosomal miRNAs examined, let-7a-5p exhibited the highest abundance in both the healthy control and IPF groups (Figure 1G).

**Fig.1.**
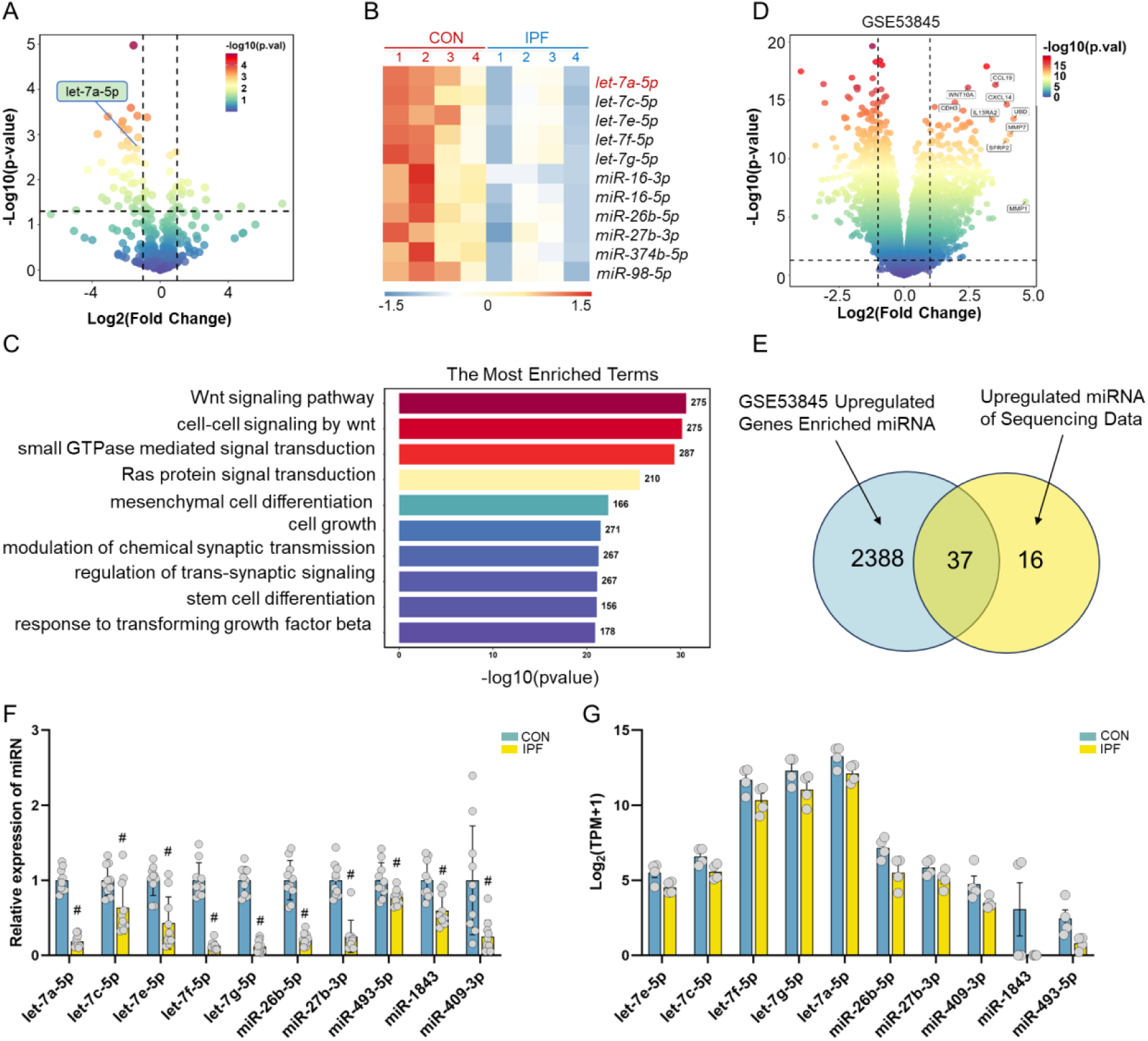
Expression of serum exosomal let-7a-5p is decreased in IPF patients.(A-B) Serum exosomal RNAs were extracted from 4 patients diagnosed with idiopathic pulmonary fibrosis and 4 healthy control subjects. miRNA microarray analysis was carried out to identify the differentially expressed miRNAs in serum exosomes. (C) The target genes of the differentially expressed miRNAs were predicted via the TargetScanHuman 8.0 website. Functional annotation of the screened differential miRNAs was conducted based on the Gene Ontology (GO) database.(D) Differentially expressed genes in the lung tissues of forty idiopathic pulmonary fibrosis patients and eight healthy controls were analyzed from the NCBI public database GSE53845 (log₂ > 1 signified up - regulation, and log₂ < 1 signified down-regulation). (E) The miRNAs recruited by up-regulated genes in lung tissue obtained from the GSE53845 dataset were intersected with the sequencing data of this study. (F) For the overlapping miRNAs, PCR validation was performed with an expanded sample size of 10 cases to confirm the sequencing results. (G) Expression abundance of the 10 miRNAs in serum exosomes in the healthy control group and IPF group. Data were presented as mean ± SEM. The independent samples t-test was employed for normally distributed data, and the Mann-Whitney U test was utilized for non-normally distributed data (*p < 0.05, #p <0.01).

### Let-7a-5p is negatively associated with pulmonary fibrosis in humans

Serum samples from patients with IPF were analyzed for the expression of serum exosomal let-7a-5p and KL-6. The expression of let-7a-5p was decreased (Figure 2A) and that of KL-6 was increased (Figure 2B), and the two showed a negative correlation (Figure 2C). Blood gas analysis, which reflects pulmonary gas exchange, revealed that let-7a-5p was positively correlated with arterial partial pressure of oxygen (PaO₂) (Figure 2D). Pulmonary function tests assessing lung ventilation demonstrated that let-7a-5p was positively correlated with forced expiratory volume in one second (FEV1), forced vital capacity (FVC), and diffusing capacity of the lungs for carbon monoxide (DLCO) (Figure 2E-G). However, the level of let-7a-5p was negatively correlated with the lung CT score (Figure 2H). Together, these data suggest that the serum level of Let-7a-5p was negatively associated with patients with pulmonary fibrosis.

**Fig.2.**
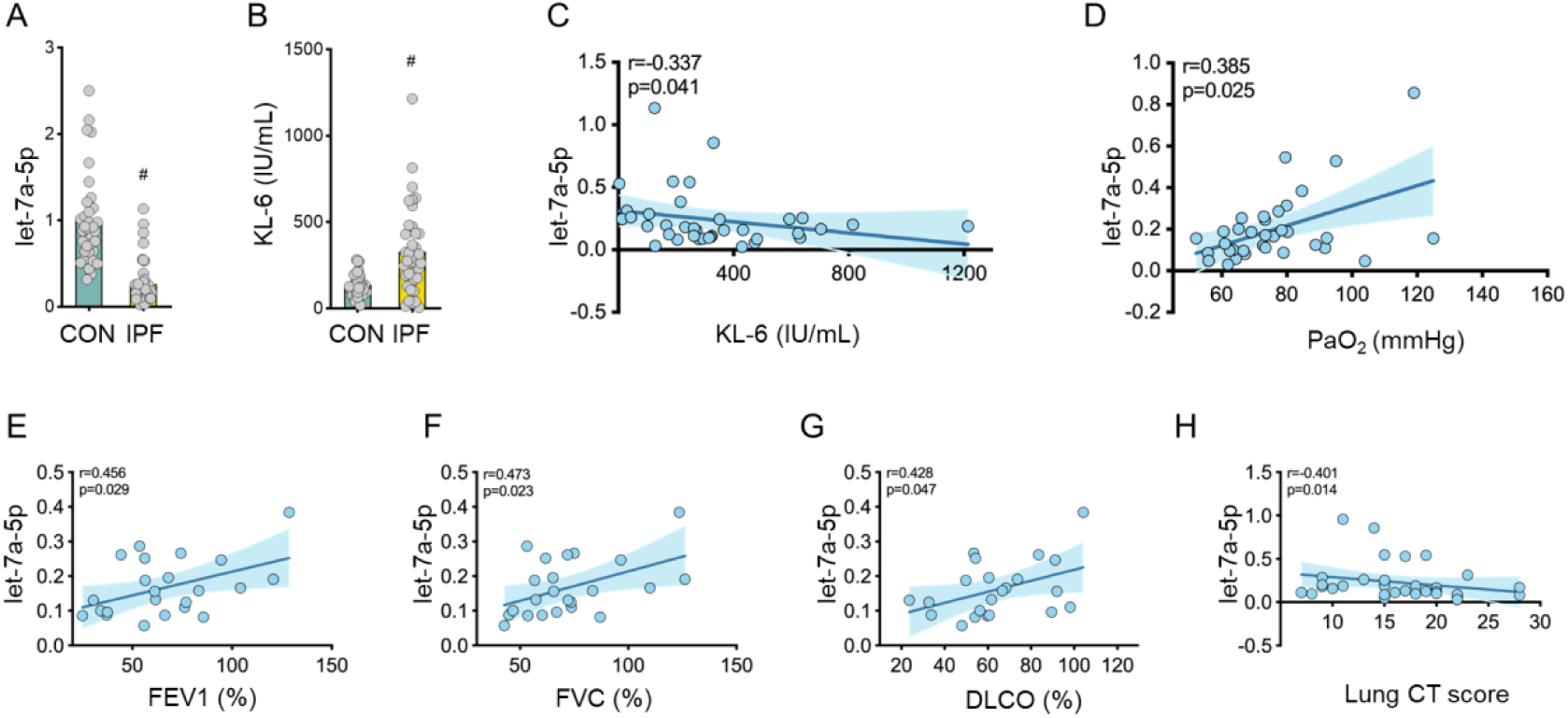
Let-7a-5p is associated with clinical characteristics of IPF patients. (A) Serum exosomal miRNA was extracted from 34 healthy controls and 43 patients with idiopathic pulmonary fibrosis, and let-7a-5p expression was detected by PCR. (B) Serum KL-6 expression was measured using an ELISA kit. (C) Correlation analysis between let-7a-5p and KL-6. (D) Correlation analysis of let-7a-5p with partial pressure of oxygen (PaO₂). (E-G) Correlation analysis of let-7a-5p with pulmonary function parameters in pulmonary fibrosis patients: FEV1, FVC and DLCO. FEV1, FVC and DLCO values are reported as measured/predicted values for each patient. (H) Correlation analysis of let-7a-5p with lung CT score in pulmonary fibrosis patients. Data are presented as mean ± SEM. The Pearson test was used for normally distributed data, and the Spearman test was used for non-normally distributed data (*p<0.05, #p <0.01; r>0 indicates a positive correlation, r<0 indicates a negative correlation).

### Let-7a-5p mediates EndoMT via regulation of TGFBR1

Given that Gene Ontology (GO) database indicated that the TGF-β signaling pathway is regulated by the differentially expressed miRNAs, we used TargetScanHuman 8.0 to identify that let-7a-5p targets the 3’ untranslated region (3’UTR) of TGFBR1 mRNA (Figure 3A).

**Fig.3.**
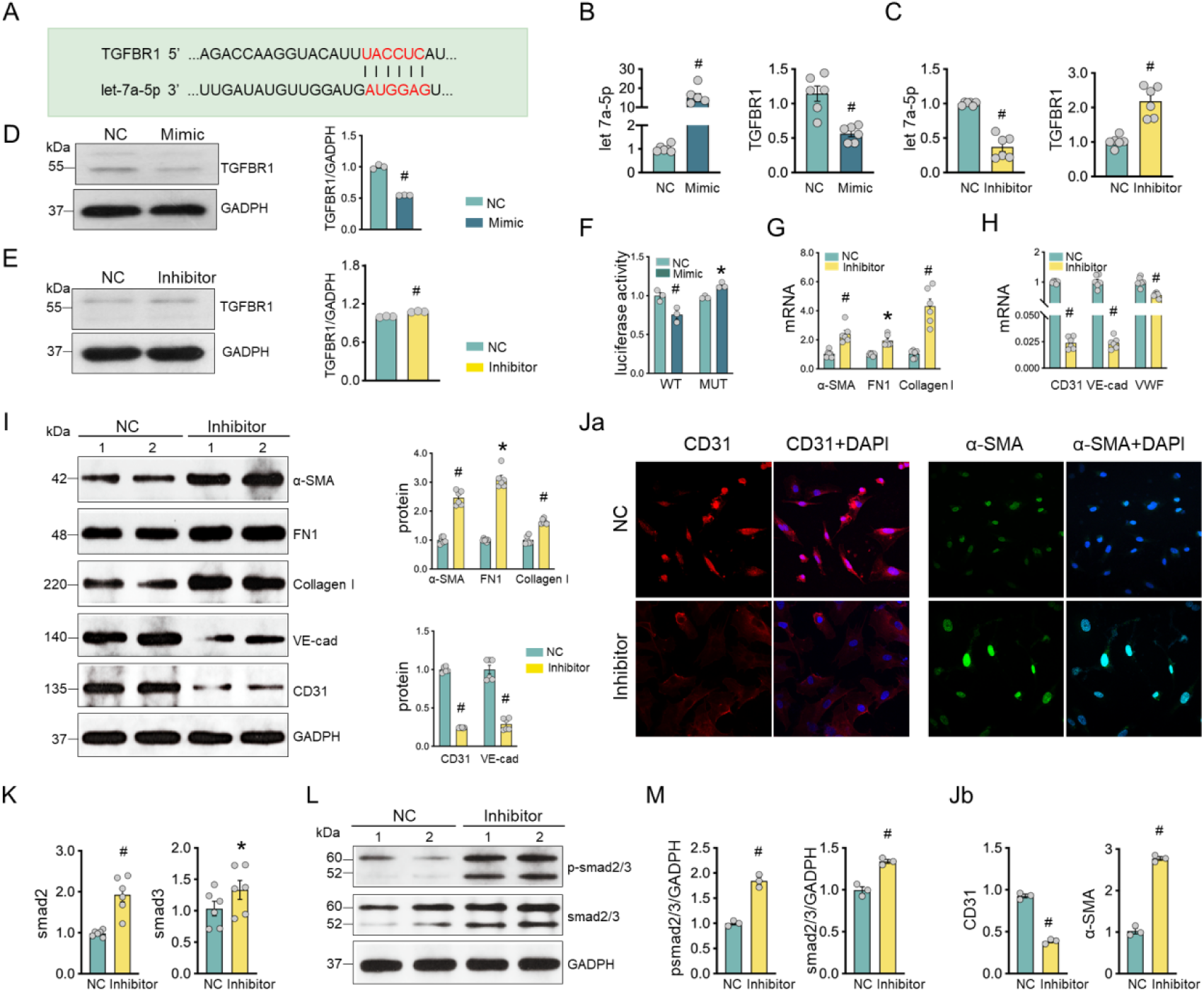
let-7a-5p mediates EndoMT via downregulation of TGFBR1. (A) Prediction of the let-7a-5p binding site on the 3’UTR of TGFBR1 using TargetScanHuman 8.0. (B) Endothelial cells were seeded in 6-well plates at a density of 1×10⁶ cells per well. Subsequently, the cells were transfected with let-7a-5p mimic using lipo2000, and the expression of the TGFBR1 gene was detected via PCR. (C) PCR-based detection of TGFBR1 gene expression in endothelial cells subsequent to let-7a-5p inhibition. (D) Western blot analysis of TGFBR1 protein expression in endothelial cells following let-7a-5p overexpression. (E) Western blot analysis of TGFBR1 protein expression in endothelial cells after let-7a-5p inhibition. (F) Wild-type and mutant TGFBR1 plasmids were designed and co-transfected with let-7a-5p mimic into 293T cells. Fluorescence intensity was measured using a dual-luciferase reporter assay kit; the binding of let-7a-5p to TGFBR1 leads to fluorescence quenching. (G-H) Endothelial cells were seeded in 6-well plates at a density of 1×10⁶ cells per well. The cells were then transfected with let-7a-5p inhibitor using lipo2000, and the gene expression of mesenchymal and endothelial markers was detected by PCR. (I) Western blot analysis of mesenchymal and endothelial marker protein expression in endothelial cells after transfection with let-7a-5p inhibitor. (Ja-Jb) Endothelial cells were cultured on coverslips in 6-well plates, transfected with let- 7a-5p inhibitor, and the expression of α-SMA and CD31 was detected by immunofluorescence. (K-M) Western blot analysis of downstream TGFβ1 pathway components (smad2/3 and p-smad2/3) in endothelial cells after transfection with let-7a-5p inhibitor. For PCR experiments, the statistical graphs present data from 6 biologically independent experiments. For western blot and immunofluorescence statistical graphs, the data represent 3 biologically independent experiments. The data are presented as mean ± SEM. For data conforming to a normal distribution, an independent samples t-test was employed; for data not conforming to a normal distribution, the Mann-Whitney U test was utilized (*p < 0.05, #p <0.01).

Functional validation demonstrated that the overexpression of let-7a-5p in ECs led to a significant downregulated TGFBR1 at both mRNA and protein levels (Figure 3B, 3D). In contrast, the inhibition of let-7a-5p resulted in a marked upregulation of TGFBR1 at both the transcriptional and translational levels (Figure 3C, 3E). We then used the dual-luciferase reporter assays to explore whether let-7a-5p directly targets the TGFBR1 3’UTR. When let-7a-5p was overexpressed in ECs, the luciferase activity was significantly reduced, and this effect was abolished by the mutation of the seed sequence targeted by let-7a-5p (Figure 3F). Functionally, let-7a-5 suppression induced a phenotypic shift characteristic of EndoMT. The expression of mesenchymal markers, including α-SMA, FN1 and collagen I, was significantly increased. Concurrently, the expression of endothelial-specific markers, such as CD31, VE-cadherin and VWF, was decreased at both the mRNA and protein levels (Figure 3G-I). Consistently, immunofluorescence staining revealed enhanced α-SMA and diminished CD31 signal in ECs with let-7a-5p inhibition by antisense oligonucleotide (ASO) (Figure 3Ja-b). Let-7a-5p suppression of TGF-β1 signaling was reinforced by the increased phosphorylation of Smad2/3 following let-7a-5p knockdown (Figure 3K-M).

### Let-7a-5p inhibition of EndoMT via TGFBR1

To test whether Let-7a-5p inhibits EndoMT, cultured ECs were firstly stimulated with varying concentrations of bleomycin. At a concentration of 25 μg/ml, the expression of CD31 exhibited the most pronounced decline, while the expression of α-SMA increased (Figure 4A), suggesting the occurrence of bleomycin to induce EndoMT. Thus, 25 μg/ml was chosen as the experimental concentration for subsequent investigations. Following bleomycin induction, the expression of let-7a-5p in ECs was diminished (Figure 4B), but that of TGFBR1 was upregulated (Figure 4C-E). Moreover, levels of mesenchymal markers (α-SMA, FN1, and collagen I) were elevated, while those of endothelial markers (CD31, VWF, and VE-cadherin) were reduced (Figure 4F-H). Importantly, let-7a-5p overexpression mitigated the bleomycin-induced TGFBR1 (Figure 4I-J) and mesenchymal markers, but restored the expression of EC markers (Figure 4K-M). Immunofluorescence assays further corroborated that overexpression of let-7a-5p suppressed the bleomycin-induced α-SMA and restored CD31 (Figure 4N-P). These findings indicate that let-7a-5p overexpression attenuated the bleomycin-induced EndoMT, possibly through suppressing TGFBR1.

**Fig.4.**
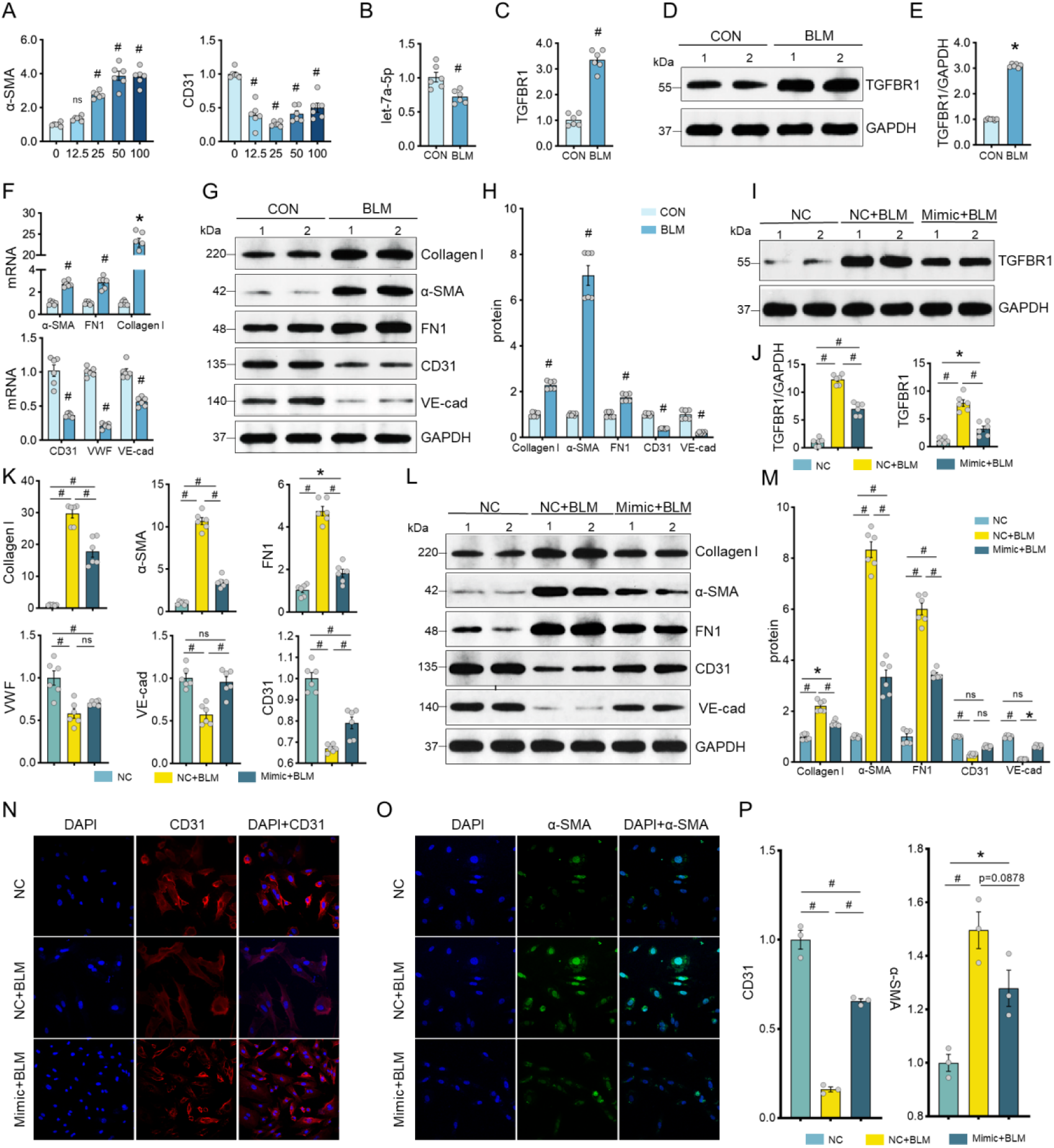
Overexpression of let-7a-5p inhibits EndoMT via TGFBR1 in vitro. (A) Endothelial cells were stimulated with bleomycin at concentrations of 0 μg/ml, 12.5 μg/ml, 25 μg/ml, 50 μg/ml, and 100 μg/ml. The expression levels of the mesenchymal marker α-SMA and the endothelial marker CD31 were detected via PCR. The concentration of 25 μg/ml was selected for subsequent experimental procedures. (B-E) Endothelial cells were seeded in 6-well plates at a density of 1×10⁶ cells per well and treated with 25 μg/ml bleomycin. The expression levels of let-7a-5p and its target gene TGFBR1 were detected using PCR and western blotting techniques. (F-H) Subsequent to stimulation with 25 μg/ml bleomycin, the mRNA and protein expression levels of mesenchymal and endothelial markers in endothelial cells were detected by means of PCR and western blotting. (I-J) Endothelial cells were transfected with let-7a-5p mimic using Lipo2000. After a 6 hour interval, the medium was replaced with a medium containing 25 μg/ml bleomycin. RNA and protein samples were harvested 24 hours later, and the expression of TGFBR1 was detected through PCR and western blotting. (K-M) Endothelial cells were treated in accordance with the protocol described in panel I. The expression levels of mesenchymal and endothelial markers were detected using PCR and western blotting. (N-P) Endothelial cells cultured on coverslips in 6-well plates were transfected with let-7a-5p mimic. After 6 hours, the medium was replaced with a medium containing 25 μg/ml bleomycin. Immunofluorescence assays were performed 24 hours later to detect the mesenchymal marker α-SMA and the endothelial marker CD31. Statistical graphs for PCR represent data obtained from 6 biologically independent experiments. Statistical graphs for western blotting and immunofluorescence represent data derived from 3 biologically independent experiments. Data are presented as the mean ± SEM. For data exhibiting a normal distribution, an independent samples t-test was employed; for non-normally distributed data, the Mann-Whitney U test was utilized. Multi-group comparisons employed one-way ANOVA (normal) or Kruskal-Wallis test (non-normal) (*p < 0.05, #p <0.01).

### let-7a-5p overexpression mitigates pulmonary fibrosis in mice

To investigate whether let-7a-5p supplementation can mitigate pulmonary fibrosis, let-7a-5p agomir was administered to the bleomycin-administered mice through tail-vein injection (Figure 5A). Body weight alterations were documented, and the mice were euthanized on day 21. The results indicated no significant disparity in body weight between the agomir NC + BLM group and the agomir + BLM treatment group. Nevertheless, both groups had significant body weight, in comparison to the agomir NC group (Figure 5B). HE staining of lung tissues demonstrated a notable amelioration of pulmonary fibrosis in the agomir + BLM treatment group (Figure 5C). Although the alveolitis score did not show significant improvement (Figure 5D), the pathological fibrosis score was significantly diminished (Figure 5E), by let-7a-5p agomir.

**Fig.5.**
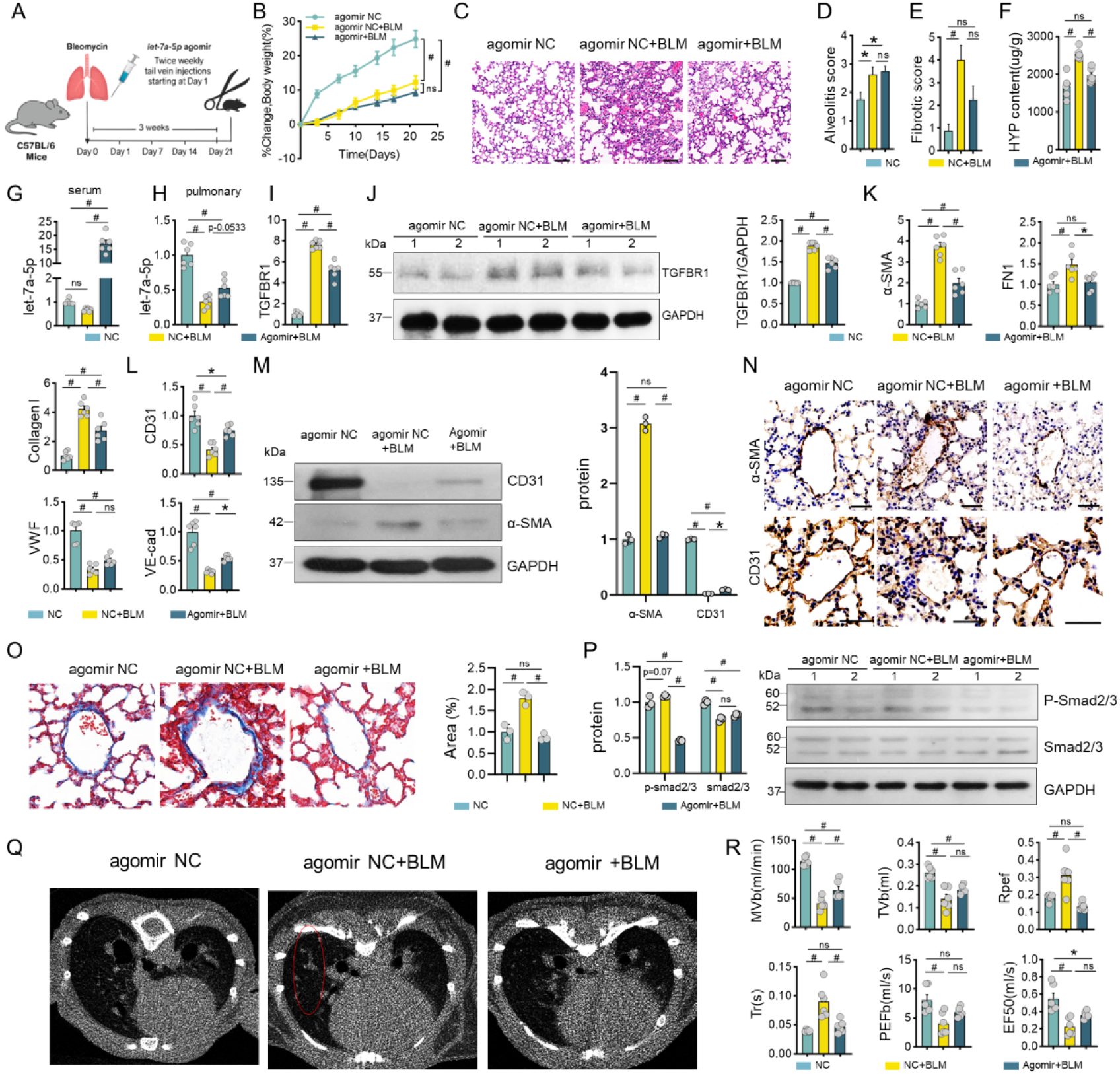
let-7a-5p inhibits pulmonary EndoMT and fibrosis in mice. (A) C57BL/6J mice were administered a single intratracheal instillation of bleomycin to establish a pulmonary fibrosis model. The day of modeling was designated as day 0. Starting from day 1, let-7a-5p agomir (15 mg/kg) was injected via the tail vein twice a week. Body weight was measured twice weekly, and mice were sacrificed on day 21. Each group consisted of 6 mice. (B) Body weight changes in the control group (agomir NC), model group (agomir NC+BLM), and treatment group (agomir+BLM). (C) Lung tissues from the control, model, and treatment groups were fixed in 4% paraformaldehyde, paraffin-embedded, and stained with HE staining. (D) Szapiel alveolitis score. (E) Ashcroft pulmonary fibrosis pathological score. (F) Hydroxyproline content in lung tissues of the control, model, and treatment groups was measured using a hydroxyproline assay kit. (G-H) Expression of let-7a-5p in serum exosomes and lung tissues of the control, model, and treatment groups was detected by PCR. (I-J) mRNA and protein expression of TGFBR1 in lung tissues of the control, model, and treatment groups was detected by PCR and Western blot. (K-M) mRNA and protein expression of mesenchymal and endothelial markers in lung tissues of the control, model, and treatment groups was detected by PCR and Western blot. (N) Immunohistochemical detection of α-SMA and CD31 expression in each group; brown color indicates positive expression. (O) Masson staining to assess collagen deposition in each group; blue color indicates collagen fibers. (P) Western blot analysis of phosphorylated smad2/3 levels in lung tissues of each group. (Q) Mice were anesthetized with isoflurane, and lung imaging was performed using a micro-CT scanner (Scanco VivaCT80, Switzerland) with scans taken at 30-μm intervals to observe pulmonary fibrosis in each group. White patches indicate fibrotic areas. (R) Pulmonary function was assessed using a non-invasive pulmonary function testing system. Statistical graphs for PCR represent data from 6 biologically independent experiments. Statistical graphs for Western blot, immunohistochemistry, and Masson staining represent data from 3 biologically independent experiments. Data are presented as mean ± SEM. For normally distributed data, an independent-samples t-test was used; for non-normally distributed data, the Mann-Whitney U test was used. Multi-group comparisons employed one-way ANOVA (normal) or Kruskal-Wallis test (non-normal) (*p < 0.05, #p <0.01).

When compared with the agomir NC+BLM group Histologically, the agomir+ BLM treatment group showed a significantly elevated level of let-7a-5p in serum exosomes and lung tissue (Figure 5G-H), but decrease in hydroxyproline content and expression of TGFBR1 in lung tissues (Figure 5F, 5I-J). In line with these results, levels of mesenchymal markers (α-SMA, FN1, collagen I) were decreased, whereas those of EC markers (CD31, VE-cadherin) was significantly increased (Figure 6K-M). Immunohistochemical findings also support such decrease in EndoMT (Figure 5N). Masson staining indicated a markedly decreased collagen deposition in the treatment group (Figure 5O). Compared with the agomir NC+ BLM group, the activation of smad2/3 was significantly suppressed in the treatment group (Figure 5P). Furthermore, pulmonary CT imaging showed a significant reduction in fibrotic shadows (Figure 5Q). Pulmonary function tests demonstrated a significant increase in minute ventilation (MVb), with non-significant upward trends in tidal volume (TVb), peak expiratory flow (PEFb), and mid-expiratory flow (EF50), alongside significant decreases in peak expiratory time ratio (Rpef) and relaxation time (Tr). Overall, these findings indicate that let-7a-5p may alleviate pulmonary fibrosis, likely through targeting the TGF-β signaling pathway (Figure 5R).

**Fig.6.**
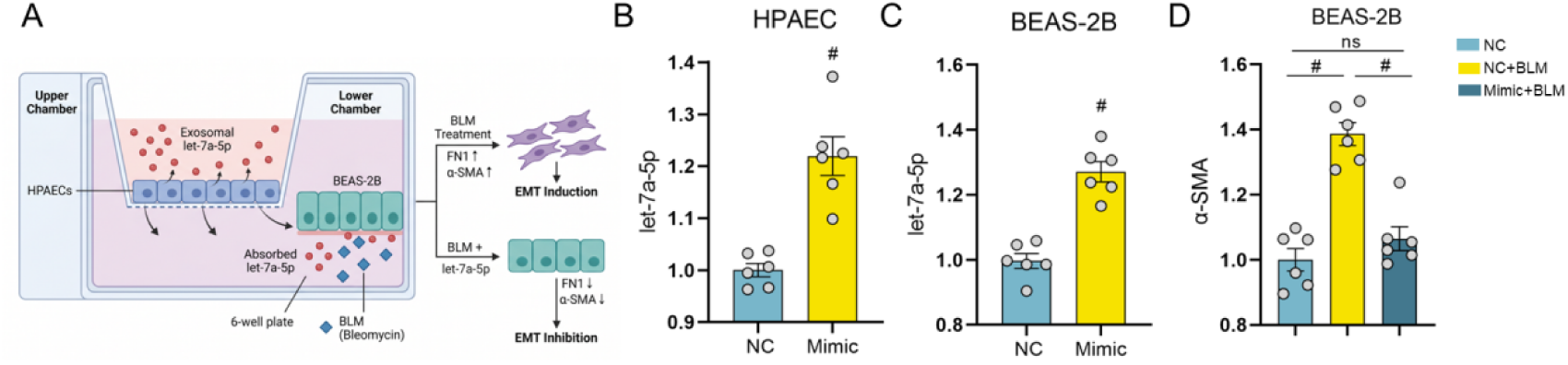
Overexpression of let-7a-5p in endothelial cells alleviates EMT.(A) Schematic representation of the co-culture system of HPAECs and BEAS-2B. (B) Overexpression of let-7a-5p in HPAECs in the upper chamber of the Transwell system. (C) let-7a-5p expression in BEAS-2B in the lower chamber of the Transwell system. (D) After treating endothelial cells in the upper chamber with NC, NC+BLM, or let-7a-5p mimic+BLM, the gene expression of α-SMA in BEAS-2B in the lower chamber was detected. For normally distributed data, an independent-samples t-test was used; for non-normally distributed data, the Mann-Whitney U test was used. Multi-group comparisons employed one-way ANOVA (normal) or Kruskal-Wallis test (non-normal) (*p < 0.05, #p <0.01).

### Overexpression of let-7a-5p in endothelial cells alleviates EMT

Using a Transwell chamber, HPAECs were co-cultured with bronchial epithelial cells (BEAS-2B), with HPAECs in the upper chamber and BEAS-2B in the lower chamber. Overexpression of let-7a-5p in HPAECs resulted in increased let-7a-5p expression in BEAS-2B (Figure 6A-C), demonstrating that endothelial-derived let-7a-5p can be secreted by endothelial cells and taken up by epithelial cells. Based on this finding, we investigated the effect of endothelial-secreted let-7a-5p on bleomycin-induced EMT. The results showed that let-7a-5p inhibited the expression of α-SMA in BEAS-2B (Figure 6D), indicating that endothelial cell-derived let-7a-5p also suppresses EMT.

## DISCUSSION

This study centers on a new pathomechanism involving let-7a-5p targetng TGFBR1. We show the downregulated let-7a-5p in pulmonary fibrotic tissues and serum, and its expression levels are closely correlated with the severity of fibrosis. The loss of let-7a-5p during the onset of pulmonary fibrosis leads to the upregulation of TGFBR1 and the activation of TGF-β pathway and induction of EndoMT. Moreover, this EndoMT process can further influence EMT. Collectively, these findings demonstrate a protective role of let-7a-5p in lung endothelium against pulmonary fibrosis, providing the first comprehensive validation of the let-7a-5p/TGFBR1/EndoMT regulatory axis in both cellular and animal models.

The pathogenesis and progression of pulmonary fibrosis is a complex process entailing the coordinated actions of several cell types and many signaling pathways^[29–31]^. Both ECs and epithelial cells play significant roles in the fibrotic process^[32, 33]^. In recent years, the unique functions of ECs have received increasing attention. Lung endothelium constitute the barrier of the pulmonary vascular, dysfunction is a key pathophysiological factor driving the initiation and progression of pulmonary fibrosis^[34, 35]^. Under normal physiological conditions, pulmonary ECs maintain an intact arrangement and stabilize pulmonary circulation by secreting vasoactive substances such as nitric oxide and prostacyclin^[36]^. However. EndoMT is a core mechanism through which healthy ECs are converted to mesenchymal-like cells, resulting in the increased synthesis and deposition of ECM^[37, 38]^. EndoMT can further promote pulmonary fibrosis through triggering inflammatory responses, secreting exosomes, and participating in vascular rarefaction and dysfunction^[39, 40]^.

Although the let-7 family has been implicated in the anti-fibrotic effects against IPF in various cell types, the specific role of let-7a-5p in regulating EndoMT via targeting TGFBR1 has not been systematically elucidated. Previous research has demonstrated that in alveolar epithelial cells, let-7a-5p can target and inhibit the expression of genes such as HMGA2 and TGF-β1, thereby reducing the EMT process and decreasing the secretion of pro-fibrotic factors, ultimately suppressing pulmonary fibrosis^[41, 42]^. In fibroblasts, let-7a-5p can target and regulate collagen-coding genes such as COL1A1 and COL3A1^[43]^, inhibiting fibroblast proliferation and activation, reducing ECM deposition, and exerting anti-fibrotic effects^[44, 45]^. Our study extends these findings by providing novel cell type evidence in endothelial cells. We demonstrate that let-7a-5p can specifically bind to the 3’UTR of TGFBR1 mRNA, inhibits TGFBR1 expression, and consequently suppresses the EndoMT in ECs, reducing the generation of interstitial-like cells and ECM deposition. Moreover, we show for the first time that endothelial-derived let-7a-5p can influence EMT in epithelial cells through paracrine signaling, revealing an intercellular crosstalk mechanism that has not been previously reported. Thus, let-7a-5p in lung ECs would be beneficial, which is consistent with the anti-fibrotic effects of let-7a-5p observed in previous studies. Additionally, the downregulation of serum exosomal let-7a-5p in pulmonary fibrosis patients correlates with disease severity, suggesting its potential as a biomarker for early diagnosis and disease assessment. This provides a novel non-invasive approach for the diagnosis of pulmonary fibrosis, and identifies TGFBR1 as a potential therapeutic target.

Although this study has elucidated the role and mechanism of the serum exosome let-7a- 5p/TGFBR1/EndMT regulatory axis in pulmonary fibrosis, several limitations exist and require further investigation. Firstly, while our clinical cohort revealed a significant correlation between serum exosomal let-7a-5p levels and disease severity, validation in larger, multi-center cohorts is needed to establish its clinical utility as a biomarker. Secondly, although we identified TGFBR1 as a direct target of let-7a-5p, we cannot exclude the possibility that other target genes may also contribute to the observed anti-fibrotic effects. Further exploration of the regulatory network of let-7a-5p is warranted. Thirdly, while our co-culture experiments suggest that endothelial-derived let-7a-5p can be taken up by epithelial cells to suppress EMT, the precise source cells of serum exosomal let-7a-5p (e.g., pulmonary ECs, epithelial cells, or immune cells) and the detailed mechanisms of exosomal secretion and transport require further clarification. Furthermore, the proposed therapeutic strategies (e.g., exosomal delivery of let-7a-5p still need to be validated for safety and efficacy in pre-clinical study involving large animal models.

## Acknowledgments

We would like to express our sincere gratitude to John Y-J. Shyy from the Division of Cardiology, Department of Medicine, University of California, for his guidance on this research. We also extend our sincere gratitude to the School of Medicine at Xi’an Jiaotong University and the School of Life Sciences at Northwestern Polytechnical University for providing the necessary experimental facilities and resource support for this study. We further thank Tibikang Biotechnology for their sequencing technology support, and we are particularly grateful to the Affiliated Chest Hospital of Northwest University for their assistance in specimen collection.

## Author contributions

CRediT: Fenqing Shang: conceptualization, project administration, supervision and funding acquisition and writing-review & editing. Hui Shen and Jianying Li: conceptualization, project administration, supervision and writing-review & editing. Juan Pang: formal analysis, investigation, methodology and writing-original draft and funding acquisition. Jiarui Shen: formal analysis, visualization, writing-original draft. Wei Yang and Zhihui Wu: data curation and methodology. Xing Gu, Yuhang Xia, Ruixuan Wang, and Yujie Cao: resources, writing-review & editing and methodology. Longzhi Wang: validation and methodology. All authors reviewed and approved the final manuscript.

## Data availability

The datasets supporting the conclusions of this study are available from the corresponding author upon reasonable request.

## Ethics approval and consent to participate

Human sample collection was conducted in accordance with the Declaration of Helsinki. Animal experiments were conducted in accordance with the National Institutes of Health Guide for the Care and Use of Laboratory Animals. Human procedures were approved by the Ethics Committee of Xi’an Chest Hospital (Approval No. R2025-012-01), and animal procedures were approved by the same committee (Approval No. A2024-001-01).

## Consent for publication

All authors approved the manuscript publication in Journal of PLOS One.

## Supplementary materials

The following supplementary materials are available with the online version of this article: Supplementary Table S1, Supplementary Table S2, and Supplementary Table S3.

## REFERENCES

[1] Lachowicz J A, Steinfort D P, Smallwood N E, Prasad J D. Advances in management of pulmonary fibrosis [J]. Internal medicine journal, 2025, 55(7): 1070–80. doi:10.1111/imj.70051

[2] Fox L, Murray B. Idiopathic pulmonary fibrosis: the role of the respiratory advanced nurse practitioner [J]. British journal of nursing (Mark Allen Publishing), 2025, 34(13): 675–82.doi:10.12968/bjon.2024.0432

[3] Ishii H, Kinoshita Y, Hamada N, Fujita M, Kushima H. Idiopathic pleuroparenchymal fibroelastosis: diagnosis and management [J]. Expert review of respiratory medicine, 2025, 19(7): 697–708. doi:10.1080/17476348.2025.2499651

[4] Ali A, Glassberg M K. Managing Acute Exacerbations in Idiopathic Pulmonary Fibrosis and Progressive Pulmonary Fibrosis in the Intensive Care Unit Setting [J]. Critical care nursing clinics of North America, 2025, 37(3): 407–19. doi:10.1016/j.cnc.2025.05.002

[5] Provenzani A, Leonardi Vinci D, Alaimo M, Di Maria S, Tuzzolino F, Floridia G, Di Stefano R, Carollo A, Callari A, Polidori P, Vitulo P. Real-world insights into safety, tolerability, and predictive factors of adverse drug reactions in treating idiopathic pulmonary fibrosis with pirfenidone and nintedanib [J]. Therapeutic advances in drug safety, 2025, 16: 20420986251341645. doi:10.1177/20420986251341645

[6] Rich R L, Peterson A, Brown J. Diagnosis and Management of Acute Exacerbation of Interstitial Lung Disease in the Intensive Care Unit [J]. AACN advanced critical care, 2025, 36(4): 374–88. doi:10.4037/aacnacc2025847

[7] Cui D, Che X, An R, Li L, Cui X, Jiang L, Jin J. Current Understanding of Pulmonary Fibrosis: Pathogenesis, Diagnosis, and Therapeutic Approaches [J]. Canadian respiratory journal, 2025, 2025: 3183241. doi:10.1155/carj/3183241

[8] Zang N, Wu Y, Li P, Liu Y, Wang S, Leng J, Zhan L, Lyu X, Pang L, Wang J. Viral Pathogens and Pulmonary Fibrosis: EMT-Driven Mechanisms and Insights From Traditional Chinese Medicine [J]. Reviews in medical virology, 2026, 36(2): e70118. doi:10.1002/rmv.70118

[9] Li W, Xie Y, Chen Z, Cao D, Wang Y. Epithelial-mesenchymal transition in pulmonary fibrosis: molecular mechanisms and emerging therapeutic strategies [J]. Frontiers in medicine, 2025, 12: 1658001. doi:10.3389/fmed.2025.1658001

[10] Gorelova A, Berman M, Al Ghouleh I. Endothelial-to-Mesenchymal Transition in Pulmonary Arterial Hypertension [J]. Antioxidants & redox signaling, 2021, 34(12): 891–914. doi:10.1089/ars.2020.8169

[11] Ortiz-Zapater E, Signes-Costa J, Montero P, Roger I. Lung Fibrosis and Fibrosis in the Lungs: Is It All about Myofibroblasts? [J]. Biomedicines, 2022, 10(6). doi:10.3390/biomedicines10061423

[12] Jimenez S A, Piera-Velazquez S. Endothelial to mesenchymal transition (EndoMT) in the pathogenesis of Systemic Sclerosis-associated pulmonary fibrosis and pulmonary arterial hypertension. Myth or reality? [J]. Matrix biology : journal of the International Society for Matrix Biology, 2016, 51: 26–36. doi:10.1016/j.matbio.2016.01.012

[13] Li N, Lin Z, Zhou Q, Chang M, Wang Y, Guan Y, Li H, Zhao Y, Liu N, Jin Y, Yao S. Metformin alleviates crystalline silica-induced pulmonary fibrosis by remodeling endothelial cells to mesenchymal transition via autophagy signaling [J]. Ecotoxicology and environmental safety, 2022, 245: 114100. doi:10.1016/j.ecoenv.2022.114100

[14] Adams T S, Schupp J C, Poli S, Ayaub E A, Neumark N, Ahangari F, Chu S G, Raby B A, Deiuliis G, Januszyk M, Duan Q, Arnett H A, Siddiqui A, Washko G R, Homer R, Yan X, Rosas I O, Kaminski N. Single-cell RNA-seq reveals ectopic and aberrant lung-resident cell populations in idiopathic pulmonary fibrosis [J]. Science advances, 2020, 6(28): eaba1983. doi:10.1126/sciadv.aba1983

[15] Li R, Yin H, Wang J, He D, Yan Q, Lu L. Dihydroartemisinin alleviates skin fibrosis and endothelial dysfunction in bleomycin-induced skin fibrosis models [J]. Clinical rheumatology, 2021, 40(10): 4269–77. doi:10.1007/s10067-021-05765-w

[16] Romano E, Rosa I, Fioretto B S, Matucci-Cerinic M, Manetti M. New Insights into Profibrotic Myofibroblast Formation in Systemic Sclerosis: When the Vascular Wall Becomes the Enemy [J]. Life (Basel, Switzerland), 2021, 11(7). doi:10.3390/life11070610

[17] Yoshimatsu Y, Watabe T. Emerging roles of inflammation-mediated endothelial-mesenchymal transition in health and disease [J]. Inflammation and regeneration, 2022, 42(1): 9. doi:10.1186/s41232-021-00186-3

[18] Samarakkody A S, Cantor A B. Opening the window for endothelial-to-hematopoietic transition [J]. Genes & development, 2021, 35(21-22): 1398–400. doi: 10.1101/gad.349056.121

[19] Ying J, Wang P, Jin X, Luo L, Lai K, Li J. TGF-β1 Mediates the EndoMt in High Glucose-Treated Human Retinal Microvascular Endothelial Cells [J]. Seminars in ophthalmology, 2024, 39(4): 312–9. doi:10.1080/08820538.2023.2300806

[20] Shi Q, Liu H, Wang H, Tang L, Di Q, Wang D. MFGE8 regulates the EndoMT of HLMECs through the BMP signaling pathway and fibrosis in acute lung injury [J]. Respiratory research, 2025, 26(1): 142. doi:10.1186/s12931-025-03215-8

[21] Docshin P, Bairqdar A, Malashicheva A. Interplay between BMP2 and Notch signaling in endothelial-mesenchymal transition: implications for cardiac fibrosis [J]. Stem cell investigation, 2023, 10: 18. doi:10.21037/sci-2023-019

[22] Kimiz-Gebologlu I, Oncel S S. Exosomes: Large-scale production, isolation, drug loading efficiency, and biodistribution and uptake [J]. Journal of controlled release : official journal of the Controlled Release Society, 2022, 347: 533–43. doi:10.1016/j.jconrel.2022.05.027

[23] Guo Z Y, Tang Y, Cheng Y C. Exosomes as Targeted Delivery Drug System: Advances in Exosome Loading, Surface Functionalization and Potential for Clinical Application [J]. Current drug delivery, 2024, 21(4): 473–87. doi:10.2174/1567201819666220613150814

[24] Tian J, Han Z, Song D, Peng Y, Xiong M, Chen Z, Duan S, Zhang L. Engineered Exosome for Drug Delivery: Recent Development and Clinical Applications [J]. International journal of nanomedicine, 2023, 18: 7923–40. doi:10.2147/IJN.S444582

[25] Seasock M J, Shafiquzzaman M, Ruiz-Echartea M E, Kanchi R S, Tran B T, Simon L M, Meyer M D, Erice P A, Lotlikar S L, Wenlock S C, Ochsner S A, Enright A, Carisey A F, Romero F, Rosas I O, King K Y, Mckenna N J, Coarfa C, Rodriguez A. Let-7 restrains an epigenetic circuit in AT2 cells to prevent fibrogenic intermediates in pulmonary fibrosis [J]. Nature communications, 2025, 16(1): 4353. doi:10.1038/s41467-025-59641-1

[26] Wang W, Sinha A, Lutter R, Yang J, Ascoli C, Sterk P J, Nemsick N K, Perkins D L, Finn P W. Analysis of Exosomal MicroRNA Dynamics in Response to Rhinovirus Challenge in a Longitudinal Case-Control Study of Asthma [J]. Viruses, 2022, 14(11). doi:10.3390/v14112444

[27] Chen S Y, Chen Y L, Li P C, Cheng T S, Chu Y S, Shen Y S, Chen H T, Tsai W N, Huang C L, Sieber M, Yeh Y C, Liu H S, Chiang C L, Chang C H, Lee A S, Tseng Y H, Lee L J, Liao H J, Yip H K, Huang C F. Engineered extracellular vesicles carrying let-7a-5p for alleviating inflammation in acute lung injury [J]. Journal of biomedical science, 2024, 31(1): 30. doi:10.1186/s12929-024-01019-4

[28] Song D, Tang X, Du J, Tao K, Li Y. Diazepam inhibits LPS-induced pyroptosis and inflammation and alleviates pulmonary fibrosis in mice by regulating the let-7a-5p/MYD88 axis [J]. PloS one, 2024, 19(6): e0305409. doi:10.1371/journal.pone.0305409

[29] Moss B J, Ryter S W, Rosas I O. Pathogenic Mechanisms Underlying Idiopathic Pulmonary Fibrosis [J]. Annual review of pathology, 2022, 17: 515–46.doi:10.1146/annurev-pathol-042320-030240

[30] Unterman A, Zhao A Y, Neumark N, Schupp J C, Ahangari F, Cosme C, Jr., Sharma P, Flint J, Stein Y, Ryu C, Ishikawa G, Sumida T S, Gomez J L, Herazo-Maya J D, Dela Cruz C S, Herzog E L, Kaminski N. Single-Cell Profiling Reveals Immune Aberrations in Progressive Idiopathic Pulmonary Fibrosis [J]. American journal of respiratory and critical care medicine, 2024, 210(4): 484–96. doi:10.1164/rccm.202306-0979OC

[31] Zhou Y, Tong Z, Zhu X, Wu C, Zhou Y, Dong Z. Deciphering the cellular and molecular landscape of pulmonary fibrosis through single-cell sequencing and machine learning [J]. Journal of translational medicine, 2025, 23(1): 3. doi:10.1186/s12967-024-06031-8

[32] Confalonieri P, Volpe M C, Jacob J, Maiocchi S, Salton F, Ruaro B, Confalonieri M, Braga L. Regeneration or Repair? The Role of Alveolar Epithelial Cells in the Pathogenesis of Idiopathic Pulmonary Fibrosis (IPF) [J]. Cells, 2022, 11(13). doi:10.3390/cells11132095

[33] Zhang X, Sha Y, Wu Y, Guan H, Yang X, Wang W, Zhang W, Liu Y, Zhu L, Li Q. Targeting endothelial cells: A novel strategy for pulmonary fibrosis treatment [J]. European journal of pharmacology, 2025, 997: 177472. doi:10.1016/j.ejphar.2025.177472

[34] Ackermann M, Werlein C, Plucinski E, Leypold S, Kühnel M P, Verleden S E, Khalil H A, Länger F, Welte T, Mentzer S J, Jonigk D D. The role of vasculature and angiogenesis in respiratory diseases [J]. Angiogenesis, 2024, 27(3): 293–310. doi:10.1007/s10456-024-09910-2

[35] Lu W, Teoh A, Waters M, Haug G, Shakeel I, Hassan I, Shahzad A M, Callerfelt A L, Piccari L, Sohal S S. Pathology of idiopathic pulmonary fibrosis with particular focus on vascular endothelium and epithelial injury and their therapeutic potential [J]. Pharmacology & therapeutics, 2025, 265: 108757. doi:10.1016/j.pharmthera.2024.108757

[36] Mzimela N, Dimba N, Sosibo A, Khathi A. Evaluating the impact of type 2 diabetes mellitus on pulmonary vascular function and the development of pulmonary fibrosis [J]. Frontiers in endocrinology, 2024, 15: 1431405. doi:10.3389/fendo.2024.1431405

[37] Zhao W, Wang L, Wang Y, Yuan H, Zhao M, Lian H, Ma S, Xu K, Li Z, Yu G. Injured Endothelial Cell: A Risk Factor for Pulmonary Fibrosis [J]. International journal of molecular sciences, 2023, 24(10). doi:10.3390/ijms24108749

[38] Fließer E, Lins T, Berg J L, Kolb M, Kwapiszewska G. The endothelium in lung fibrosis: a core signaling hub in disease pathogenesis? [J]. American journal of physiology Cell physiology, 2023, 325(1): C2–c16. doi:10.1152/ajpcell.00097.2023

[39] Caporarello N, Mehta D, Tsukasaki Y, Sarkar A, Crawford B C, Bauer N R. Endothelial cell interactions with immune cells and fibroblasts in the pulmonary microenvironment: from the developing to the aging lung. Scientific session III - reSPIRE 2025 [J]. American journal of physiology Lung cellular and molecular physiology, 2025, 329(5): L667–l76. doi:10.1152/ajplung.00311.2025

[40] Rayner S G, Hung C F, Liles W C, Altemeier W A. Lung pericytes as mediators of inflammation [J]. American journal of physiology Lung cellular and molecular physiology, 2023, 325(1): L1–l8. doi:10.1152/ajplung.00354.2022

[41] Wang Y, Han T, Guo R, Song P, Liu Y, Wu Z, Ai J, Shen C. Micro-RNA let-7a-5p Derived From Mesenchymal Stem Cell-Derived Extracellular Vesicles Promotes the Regrowth of Neurons in Spinal-Cord-Injured Rats by Targeting the HMGA2/SMAD2 Axis [J]. Frontiers in molecular neuroscience, 2022, 15: 850364. doi:10.3389/fnmol.2022.850364

[42] Chen Z, Qiu J, Gao Y, Lu Q, Lin Y, Shi H. Study on the mechanism of let-7a-5p in regulating the proliferation in cervical cancer cells [J]. Clinical & translational oncology : official publication of the Federation of Spanish Oncology Societies and of the National Cancer Institute of Mexico, 2022, 24(8): 1631–42. doi:10.1007/s12094-022-02810-1

[43] Alrashed M M, Alshehry A S, Ahmad M, He J, Wang Y, Xu Y. miRNA Let-7a-5p targets RNA KCNQ1OT1 and Participates in Osteoblast Differentiation to Improve the Development of Osteoporosis [J]. Biochemical genetics, 2022, 60(1): 370–81. doi:10.1007/s10528-021-10105-3

[44] Wang L, Li T, Ma X, Li Y, Li Z, Li Z, Yu N, Huang J, Han Q, Long X. Exosomes from human adipose-derived mesenchymal stem cells attenuate localized scleroderma fibrosis by the let-7a-5p/TGF-βR1/Smad axis [J]. Journal of dermatological science, 2023, 112(1): 31–8. doi:10.1016/j.jdermsci.2023.09.001

[45] Luo Z, Sun Y, Qi B, Lin J, Chen Y, Xu Y, Chen J. Human bone marrow mesenchymal stem cell-derived extracellular vesicles inhibit shoulder stiffness via let-7a/Tgfbr1 axis [J]. Bioactive materials, 2022, 17: 344–59. doi:10.1016/j.bioactmat.2022.01.016

